# Biological Footprint of Artificial Light at Night in Rural and Developing Areas

**DOI:** 10.64898/2026.07.13.737561

**Authors:** Justin G. Boyles, Benjamin J. Merritt, Erin Koen, Corneile Minnaar

**Affiliations:** Environmental Solutions & Innovations, Inc. 4525 Este Avenue, Cincinnati, OH 45232, USA; Bradenton, FL 34212, USA; Department of Zoology, University of Cambridge, Cambridge, CB2 3EJ, UK

**Keywords:** Attenuation, Biological footprint of light, Lighting design, Light pollution, Rural lighting

## Abstract

**Context:** Artificial light at night (ALAN) has profound impacts on individual organisms and entire communities. Still, humans tend to underestimate the true biological (spatial) footprint of ALAN, in part because of our limited sensitivity to light compared to other organisms.

**Objectives:** We sought to demonstrate how far ALAN can reach into dark spaces at levels that can impact organismal behavior and physiology using a fundamental physical law, the inverse square law.

**Methods:** We created a spatially explicit model of light spread on real landscapes, parameterized using increasingly available landscape-scale vegetation data to account for attenuation through forests and blocking by topographic relief.

**Results:** Lighting types common in rural areas can produce biologically important effects more than 1 kilometer from the source, and effects of large lights might stretch 3 kilometers or more. The footprint of a light is determined by the complex and multidimensional interaction between characteristics of the light itself and the environment. For example, attenuation through a dense forest might decrease the footprint of a light more than 90% compared to the same light on a grassland. In complex environments, even small changes in light placement and characteristics can lead to large changes in the biological footprint of the light.

**Conclusions:** Designers and land stewards must account for lighting type, brightness, directionality, and reflected light to create ecologically responsible lighting. Vertical vegetation and topography strongly influence the propagation of biologically detrimental light, and environmental context is vital when planning and installing lights to minimize the biological impacts.

## Introduction

Artificial light at night (ALAN) is a ubiquitous and cosmopolitan stressor, affecting organisms, biological communities, and ecosystems worldwide (Falcón et al. 2020; Gaston et al. 2021; Knop et al. 2017; Longcore and Rich 2004). While effects of ALAN have been subject of intense study, less effort has been focused on understanding reach of light into natural environments, including the biological (spatial) footprint over which light can affect plants and animals (Gaston et al. 2021). To date, most empirical work on the biological footprint of light has focused on attraction to lights (e.g., Rodríguez et al. 2017; Rodríguez et al. 2015; Truxa and Fiedler 2012). Distances at which ALAN can have other impacts have been less thoroughly detailed, but even dim artificial light can significantly impact physiology, behavior, and community interactions (Aulsebrook et al. 2022). Understanding the reach of light into natural environments is important in many ecological contexts and for conservation planning and mitigation efforts.

Propagation of light into the environment is dictated by characteristics of the light source, the physics of light, environmental conditions, the landscape, and even the curvature of Earth (Aubé 2015; Massetti 2020). Some of these effects, like the physics of light, are well understood and straightforward to model, at least in theory. Others, like how light filters through a real environment, are more complicated. Light propagates via line-of-sight transmission but can be blocked by solid objects, reflected off natural and anthropogenic surfaces, and scattered in complex environments and the atmosphere (Aubé 2015; Luginbuhl et al. 2009). The impact of any individual light source on the natural environment is further affected by other lights in the landscape, humidity, smog, and cloud cover on a given night (Kocifaj 2011; Kyba et al. 2011; Wallner and Kocifaj 2023). In real-world applications, estimating the distance of light propagation is therefore highly context dependent, and assigning that light propagation biological significance is even more difficult.

Much of the research on the biological footprint of ALAN has focused on urban lighting and skyglow and their potentially dramatic effects on species and biological communities (e.g., La Sorte et al. 2017; Škvareninová et al. 2017). However, many important ecological impacts of ALAN are associated with lighting in rural and developing regions around the world (Koen et al. 2018; Sánchez de Miguel et al. 2021). Lights in rural areas may differ from those in nearby cities for a variety of reasons, including rates of technology adoption and differences in availability of reliable electricity (Coetzee et al. 2023; Khayam et al. 2023). Rural lights exist across a continuum of types and purposes ranging from house or farm lights, lighting on road and utility infrastructure, security lighting, construction equipment, and landscape lighting. Rural landscapes are exposed to highly directional lights, such as vehicle headlights or flashlights (i.e., torches), as well as diffuse lights, like unshielded streetlights or advertising lights, used to illuminate a broad area by emitting light in all directions (Team Nachtlichter et al. 2025). Dozens of experiments using even small artificial lighting installations like those commonly found in rural and developing areas have demonstrated resultant changes in plant growth and phenology and animal space use, distribution, behavior, and predator-prey interactions in a variety of taxa (e.g., Cesarz et al. 2023; Cravens and Boyles 2019; Holzhauer et al. 2015; Minnaar et al. 2015; Rotics et al. 2011).

Humans underestimate the biological footprint of ALAN on the landscape, in part because of how we perceive light. All animals are subject to a trade-off between visual acuity and light sensitivity (Caves et al. 2018); humans are particularly insensitive to light at low levels. For example, the American National Standards Institute (ANSI) uses a standard illuminance intensity (*E*) of 0.25 lux in calculations of light throw, approximating a bright full moon under which humans could reasonably be expected to identify objects. While not directly analogous, at least some animals are affected by illuminance intensities at or below 0.006 lux (Fig. 1) (Aulsebrook et al. 2022), comparable to starlight on a dark night. Illumination levels between 0.006 and 0.25 lux cause shifts in cortisol and melatonin (de Jong et al. 2016; Gutman et al. 2011; Kupprat et al. 2020) and change foraging behavior in vertebrates and invertebrates (Elangovan and Marimuthu 2001; Kerfoot 1967). Similar light intensities can influence plant physiology, phenology, and community structure (Bennie et al. 2016; Bucher et al. 2023).

**Figure 1.**
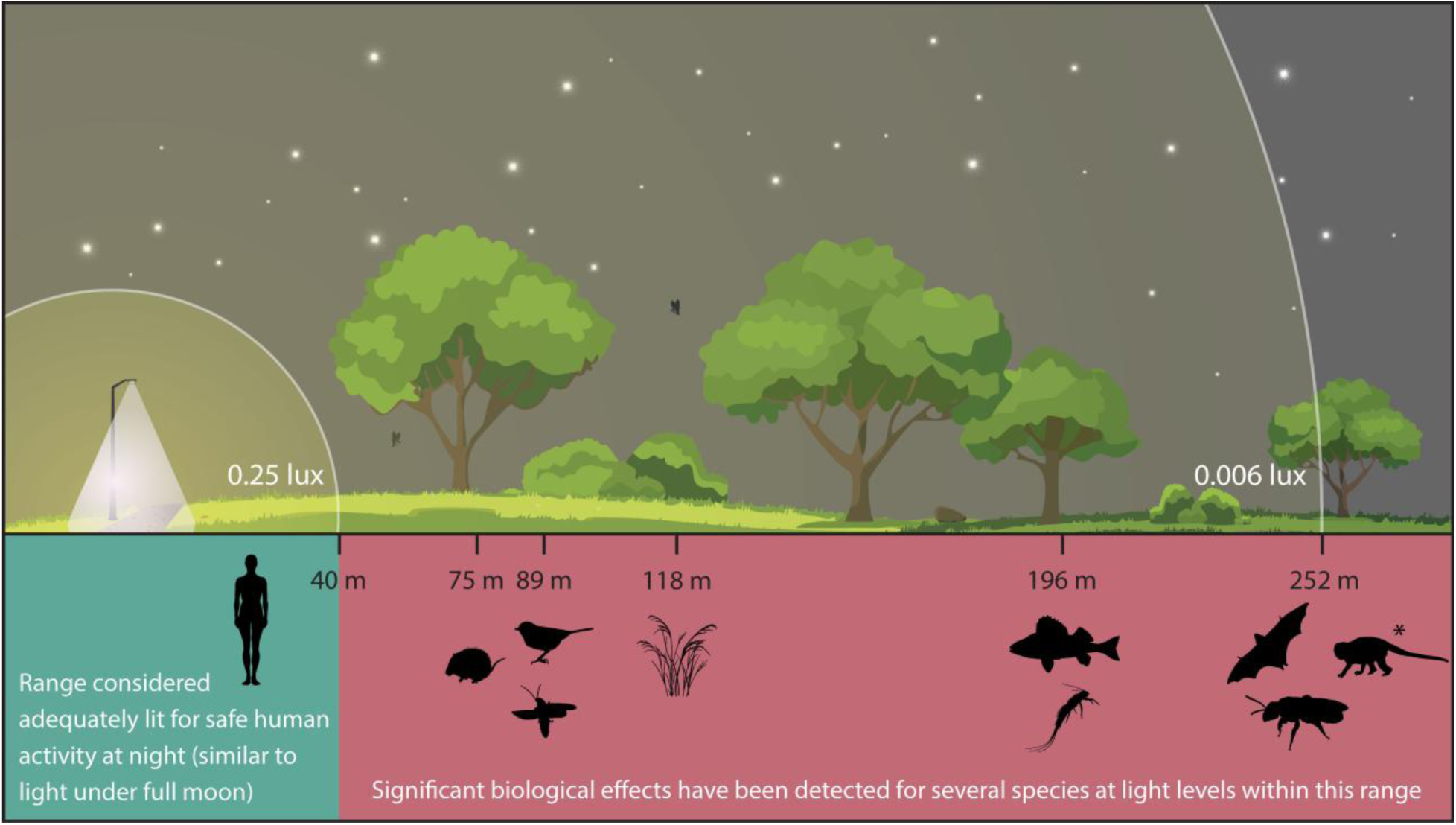
Potential distance of biological effects from a single rural light source (assuming a 5,000-lumen light source placed 5 m high, pointed downwards with a 150° beam angle). Humans can complete gross motor tasks down to ∼0.25 lux, similar to light under a full moon. Significant biological effects have been measured for species under much dimmer light. From left to right, increased cortisol and decreased foraging by spiny mice (*Acomys cahirinus* and *A. russatus*) (Gutman et al. 2011); decreased melatonin production in great tits (*Parus major*) (de Jong et al. 2016); absence of abdominal flashing activity in fireflies (*Aspisoma lineatum, A. physonotum, Pyrogaster moestus*) (Hagen et al. 2015); reduced biomass and species richness in grassland communities (16 species) (Bennie et al. 2014; Bucher et al. 2023); decreased melatonin production in Eurasian perch (*Perca fluviatilis*) (Kupprat et al. 2020); reduced entry into stream drift by various species of aquatic insects (Bishop 1969); decreased foraging by fruit bats (*Cynopterus sphinx*) (Elangovan and Marimuthu 2001); and decreased foraging activity by nocturnal bees (*Lasioglossum texanum*) (Kerfoot 1967). *At 600 m, owl monkeys (*Aotus azarai*) would experience enough illuminance to affect activity (Fernández-Duque et al. 2010). Illustration by C. Minnaar.

As light at levels that affect many organisms is imperceivable to human observers without special equipment, estimating the biological footprint of effects of ALAN beyond the human *umwelt* is a challenge. Consider a single 60-watt incandescent light bulb that produces approximately 800 lumens. That lightbulb, unshielded, would illuminate a surface to 0.25 lux at 16 m (Figure S-1). The same lightbulb would illuminate the surface to 0.006 lux at 100 m. This alone begins to frame the limitations of human perception in understanding the biological footprint of a light, but it becomes even clearer when considering how lighting technologies are applied in the real world.

## Estimating the Reach of Individual Lights into the Environment

The footprint of point-source lighting common in rural areas is difficult to quantify with relatively coarse (often 1 km^2^ grid resolution) night-time satellite imagery that has been vital in quantifying urban lighting and skyglow and its growth over the last four decades (Elvidge et al. 2014; Kyba et al. 2017; Sánchez de Miguel et al. 2021). There are many other methods and algorithms, varying in realism and complexity, to estimate the physical limits of illumination from a specified light. Modern light pollution modelling can now account for a variety of factors that influence light reach into the environment, but these models are often complex and require detailed parameterization (e.g., Bará et al. 2023). As such, they can be difficult to apply to biological questions at ecologically relevant scales (but see Bennie et al. 2014; Morrell et al. 2026; Simoneau et al. 2021). The distance at which a light maintains biologically meaningful intensity can also be roughly estimated using the inverse square law, which is a fundamental physical equation based on first principles. The inverse square law states that a quantity (in this case light) decreases at a predictable rate as the area covered increases along a spherical surface. In practice, this means the intensity of light decreases by a factor of four as distance from the source doubles. While it does not capture the importance of lighting spectrum and other characteristics (Morrell et al. 2026; van Zyl et al. 2026), the inverse square law is useful for understanding the scale of light pollution from individual lights to address ecological and conservation questions. We contend that many biologists and land stewards underestimate the full extent of biologically meaningful light pollution, so we used calculations based on the inverse square law to demonstrate the potential unseen footprint of ALAN in rural or natural areas. Calculations are contextualized in the following sections and detailed in the Supplementary Materials.

### Light Intensity and Directionality

Light intensity is the most obvious and easily understood driver of the biological footprint of a light. Brighter lights reach deeper into the surrounding environment, but the predicted relationship is not linear. A 1,000-lumen light might illuminate a surface to 0.006 lux at approximately 112 m, a 10,000-lumen light at 335 m, and 100,000-lumen light at 916 m. Directionality of a light slightly complicates the relationship but is still easily calculated in an inverse square framework (Fig. 2). Directional and shielded lighting is common in many real-world applications, and throw is determined by both the intensity of the light and the angle of the beam (i.e., how focused the light is). Modern vehicle headlights vary widely, but most are less than 5,000 lumens with a beam angle of 20-40°. Large, focused lights, like those often used in road construction, can easily surpass 100,000 lumens with beam angles of 30° or less. Some especially large directional lights can reach over 1 million lumens, but those are usually associated with airports, concert venues, sporting events, etc., and are rare in rural and developing areas. Area lights can range from less than 1,000 lumens for residential or farm lights to 50,000 lumens or more for stadium or parking lot lighting. Although lights at the high end of that range are rare in rural environments, they can be found around infrastructure (e.g., high-voltage transmission substations) or roadside services (e.g., fueling stations or rest areas).

**Figure 2.**
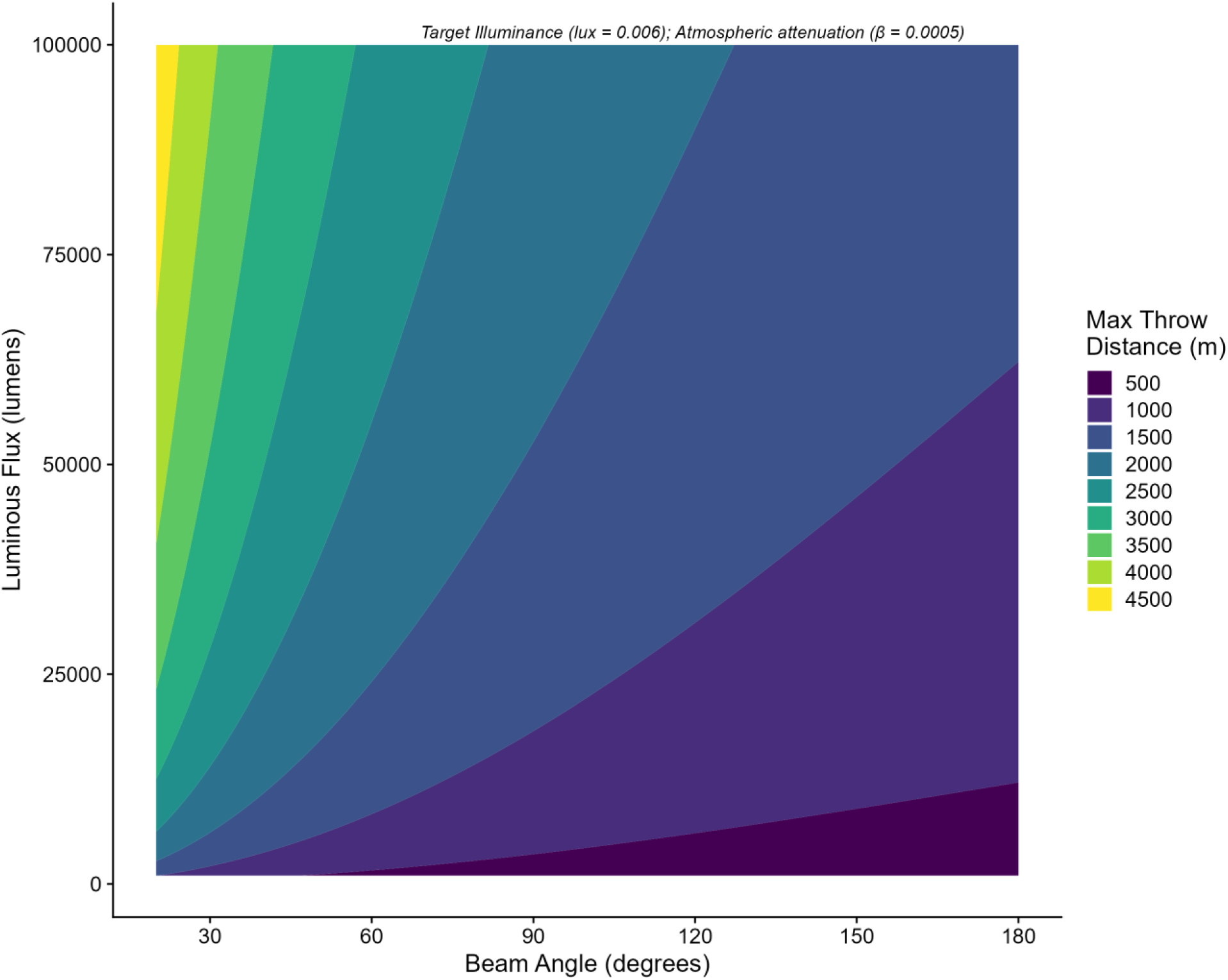
The maximum throw distance (in meters) at which a light with a given beam angle and brightness can illuminate an object to 0.006 lux, which approximates starlight. Directional lights can be circular (e.g., area lighting) or concentrated to produce a rectangular pattern (e.g., vehicle headlights and streetlighting; Flannagan 2019). Regardless of the exact pattern, directional beams are most intense at the center, with intensity falling off around the edges. For simplicity, we assume a circular beam and ignore the fall off around the edges of the beam. Values are grouped into 500 m bins for visual clarity.

In the worst-case scenario of a highly directional light aimed upwards (e.g., at a music concert) or horizontally across the landscape (e.g., a searchlight), impacts might extend 3 km or more into the environment (Fig. 2). Sustained lighting of that magnitude is probably rare in rural environments but is not impossible. A modern vehicle headlight might reach 2 km or more into the environment, although vegetation will limit that reach in many real-world scenarios (Pocock and Lawrence 2005). A bright, handheld flashlight (e.g., 1,000-lumen, 20° beam angle) could have impacts at 1 km or more (Fig. 2).

### Reflected Light, Multiple Lights, and Atmospheric Effects

Even when a lighting installation is well designed and responsibly installed, like a shielded security light directed straight down, reflected light can still reach a significant distance into the environment and have biologically important effects (Fig. 3). While more difficult to define than direct throw, the distance at which reflected light can have biological impacts can be approximated with some simplifying assumptions (e.g., the ground and anthropogenic structures act as Lambertian [i.e., diffuse] reflectors with light reflecting equally in all directions and the entire illuminated area is visible to the observer). In many real-world scenarios such assumptions will be violated, especially on natural or uneven surfaces, or for small terrestrial species at ground level. Realistically, these assumptions mean resulting estimates might be most relevant to volant nocturnal animals like insects, bats, and birds, but they can add context in other scenarios as well. The distance at which light, including reflected light, might have biological impacts is further dependent on visibility. Most rural lighting types are unlikely to reflect more than 500 m, even under high visibility conditions, but high intensity lighting (e.g., construction lighting), might reflect 1 km or more (Fig. 3). Under poor air quality conditions with low visibility, light attenuates below biologically relevant levels quickly, often in less than 100 m (Fig. 3). Conversely, these estimates extend greatly if the reflector has high albedo, for instance after a fresh snowfall or in areas with large unfinished metal surfaces.

**Figure 3.**
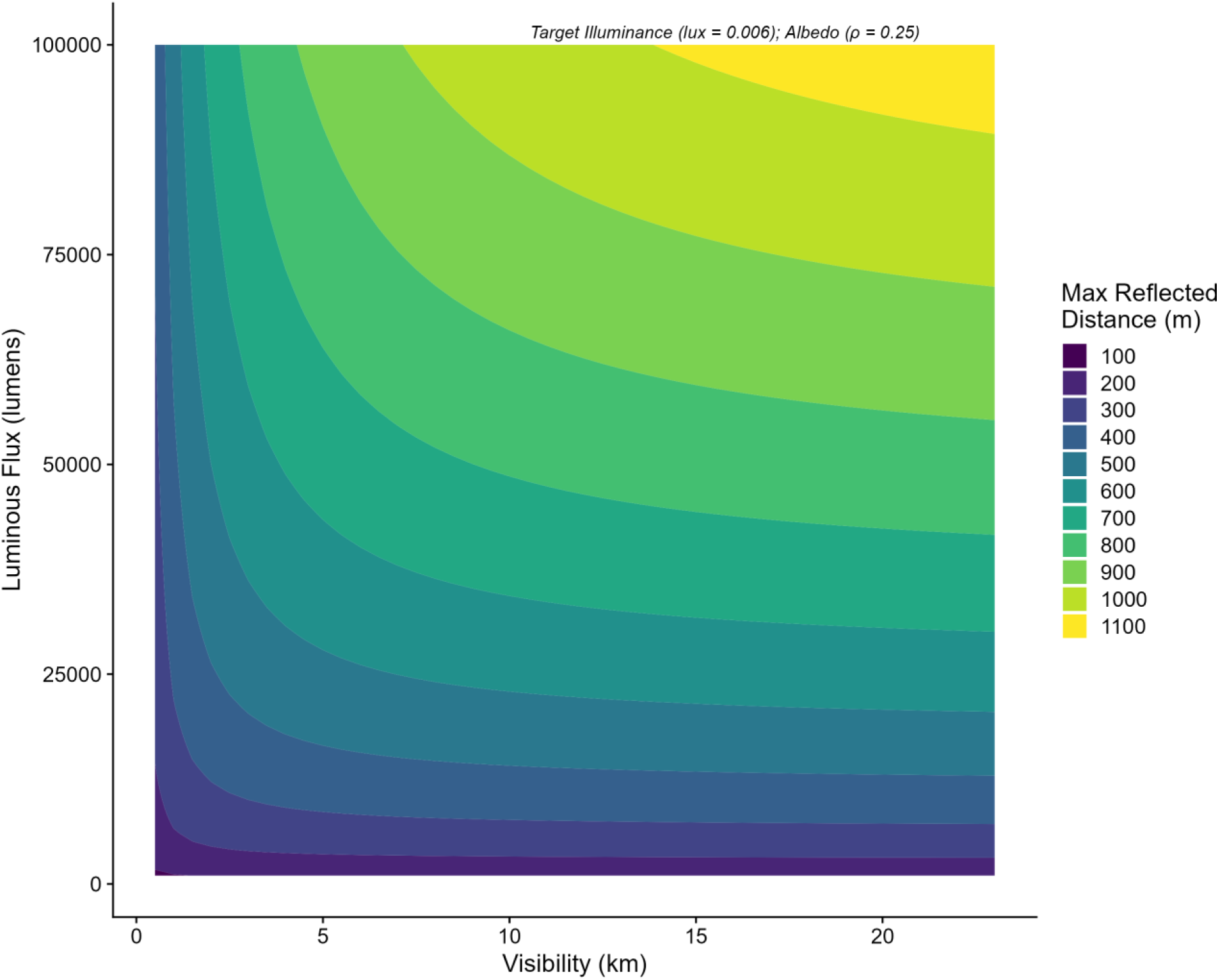
Maximum reflected distance (in meters) at which a light might be expected to illuminate an object to 0.006 lux, which approximates starlight. This simulation assumes an albedo of 0.25 and a target illuminance (*E*) of 0.006. Asphalt (tarmac) is closer to *ρ =* 0.1, while light-colored cars or unfinished metal can have *ρ >* 0.5. Fresh snow might have *ρ >* 0.8. Values are grouped into 100 m bins for visual clarity.

In many settings, multiple lights are used in an array to increase coverage, especially when used for security. When poorly designed, these large light arrays can be seen for 10 km or more^1^ and likely have detectable ecological impacts. Even when well designed, the reach into the environment can be substantial. Imagine a car park or vehicle refueling station with shielded lights facing downwards such that most light is reflected off the ground or other surfaces. In these scenarios, biologically relevant light pollution can reflect 2-3 km into the environment (Fig. S-2), rivalling the reach of large directional spotlights. Assuming that lights are positioned with some overlap to avoid dark areas, each additional light increases the impacted distance, but the relationship is not linear and less than one-to-one (Fig. S-2). An important corollary of this effect is that one light in a previously unlit area has a larger impact on the biological footprint of ALAN than the equivalent light placed in an area already saturated with light.

## Light Throw on the Landscape

Calculations based on the inverse square law are useful for framing the general scope of light pollution associated with lighting installations, but the real-world footprint of ALAN depends on both characteristics of the light and environmental context in which it is placed. Conceptually, it is easy to recognize the biological footprint of a light is greater in environments with limited topographic relief and short vegetation, like flat grasslands, deserts, or marshes, than in more complex environments where light is attenuated or blocked by vegetation, topography, and anthropogenic structures. Quantifying the biological footprint of a light in real landscapes is less intuitive because of the complex and multidimensional nature of light attenuation and shadowing.

Mapping real-world light propagation, while accounting for attenuation through vegetation and blocking of light by topographic relief (e.g., hill shadowing), is a natural extension of the inverse square model made possible by increasing availability of detailed, high-resolution data on forest structure and topography. For example, the LANDFIRE series of datasets (www.landfire.gov) provides 30 m resolution mapping of vegetation structure across the conterminous United States. Light attenuates in vegetation based on forest height, canopy cover, and canopy density, and is blocked when topographic relief rises or falls, creating a shadow. LANDFIRE products are available for each of these factors (LANDFIRE 2024a, b, c, d), allowing for approximation of the biological footprint of ALAN under different environmental conditions.

We built a simple model of light emanating from a focal grid cell then interacting with each progressive cell across a landscape described by LANDFIRE data (see Supplementary Materials for details). Light is assumed to propagate to each cell on the landscape along rays calculated with Bresenham’s algorithm (Bresenham 1965), which determines the most efficient method to move from one pixel to another in gridded systems. Upon reaching each cell along a ray, light attenuates based on canopy cover and density of the forest (if the trees are tall enough to intersect the angle of the light emanating from the focal grid) and is potentially shadowed by topography. Each cell is assigned a resulting light intensity value (in lux), with spread terminating along a ray when the resultant light intensity drops below the value assumed to have biological effects (0.006 lux in this parameterization).

The model provides broader perspective of propagation across the landscape (Fig. 4; see additional examples in Fig. S-3), allowing for further evaluation of discrete light sources and their effects. In flat areas with limited tree cover, light propagates unimpeded and the inverse square law provides a reasonable estimate of the biological footprint of a light (Fig. 4, Inset 1). Shadowing causes sometimes unpredictable and spatially heterogeneous patterns of light, especially in topographically complex areas characterized by hills and valleys (Fig. 4, Insets 2 and 3). Topography and vegetation further interact to limit the biological footprint of a light in areas with both moderate topographic relief and forest cover (Fig. 4, Inset 3). In vertically structured habitats like dense forest, vegetation likely constrains the biological footprint of the light more than topography (Fig. 4, Inset 4). Large topographic features can also be important, but slope alone is insufficient to block light as modelled here (i.e., omnidirectional spread) without small-scale undulations. Slope will be more important if the model is conceptualized for directional or shielded lighting where the light is focused downwards.

**Figure 4.**
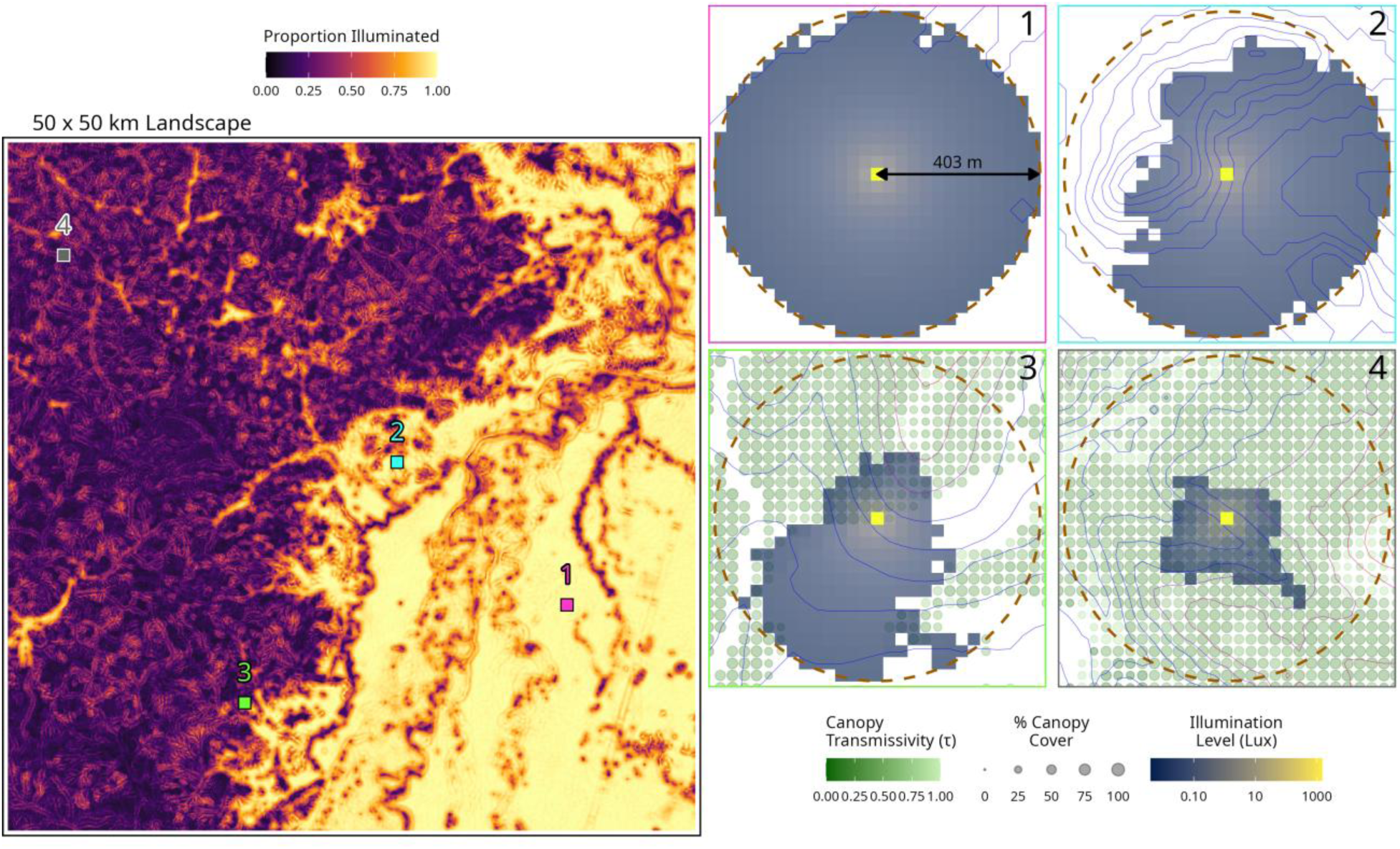
Model derived light propagation emanating from a 15,000-lumen, unshielded light atop a 10 m pole, which could represent a large light typical of roadway or infrastructure lighting. Simple modifications can be made to the model to include directional lighting, change the intensity or height of the light, or change the sensitivity of the receiver (e.g., increasing target lux to 0.25 for questions related to human safety). **Left**: a 50 x 50 km landscape detailing variation in the overall footprint of the light if placed in each 30 x 30 m grid cell, relative to the same light spreading in a treeless, flat landscape. High illumination proportions indicate a large biological footprint. Low levels indicate the light is heavily attenuated by forest or blocked by topographic relief. **Right**: Examples of light propagation for points on the larger landscapes. The brown dashed halo represents the theoretical maximum distance at which this light (bright yellow cell in the center) could illuminate the landscape to 0.006 lux based on the inverse square law. The illumination level represents every cell that is illuminated to that level under the modelled lighting conditions. Red contour lines are areas at higher elevation than the light (including the height of the pole). Blue contour lines are areas at lower elevation than the light.

What is perhaps most surprising is the scale of variation in the biological footprint of a light depending on the environmental context in which it is placed. For example, in tall, dense forests, attenuation alone can decrease the footprint of a light by 90% or more (Fig. 4, Inset 4; Fig. S-3A), but the rate of attenuation varies across forest type and even forest management strategies (Fig. S-3B; D). Topographic shadowing can have similar or greater effects (Fig. S-3C; E), but depends on the relative height of the light and undulations in the landscape. Importantly, both topography and vegetation create patterns that are unforeseeable and unpredictable to a human observer at ground level. Small-scale shifts in light placement, even less than 100 m, can lead to large relative changes in the biological footprint of light, especially in complex habitats.

Extending the inverse square law to map spread through real environments creates a semi-realistic, but scalable model capable of capturing variation in the biological footprint of a light placed in different locations on a landscape. This represents a trade-off between realism and practicality, capable of producing both detailed maps of the potential biological footprint of an individual light placed at any point on the landscape and landscape-scale estimates of the relative footprint of the light. While not meant to replace or compete with more detailed and complex models of light pollution, such models can be applied at scale and provide useful information for minimizing the biological footprint of individual lights and for large-scale deployment of lighting across the landscape.

## Managing the Biological Footprint of Artificial Light at Night

There have been concerted efforts to limit the impact of ALAN, even as lights on the landscape have increased around the world. Most of this effort has been directed at understanding and mitigating the strongest impacts of ALAN near the light source by modifying lighting color or type, shielding lights to avoid upward glow, and using technologies like motion detection that limit service times. Less effort has been expended to limit the footprint of ALAN through modifications of lighting technologies (but see Morrell et al. 2026). Some current efforts might even have the opposite effect. For example, shielding lights can, if done poorly, act to focus the light, potentially increasing the reflective distance into the atmosphere where it negatively affects nocturnal birds, bats, and insects.

Lighting footprint should be considered along with light color, type, and service time when designing lighting to minimize biological impacts. This requires a more thorough consideration of light propagation into the environment and requires further recognition of effects of ALAN that are beyond the human *umwelt*. While light propagation is more context dependent than light color or service times, simple decisions, like using the dimmest possible light to accomplish the goal of the lighting installation, decreasing the height of the lighting installation, and considering the albedo of the illuminated surface can greatly reduce the biological footprint of lights in rural and developing areas. Such decisions take on added importance in environments with limited vertical vegetation and topographic relief, where light can propagate unimpeded deep into the dark environment.

## Conclusions

Among the most important steps in managing the impacts of ALAN is establishing a broader recognition that biological footprints of even relatively small lighting installations can reach far beyond the area perceived by humans and deep into the surrounding environment. Establishing such recognition is most feasible in professional circles of planners, engineers, land stewards, and conservationists who can directly evaluate the characteristics of a light and the environmental context in which it is placed, then act to minimize ALAN and its biological impacts while accomplishing the safety or aesthetic purposes of the light. However, most lights around the world are installed without professional planning, and building a broader recognition that lights have impacts far beyond the area perceived by the human eye will be challenging. At least attempting to do so is especially important in rural and developing areas where lighting installations are increasing at the interface of anthropogenic and natural environments, but consideration of the true reach of ALAN into the environment is necessary everywhere.

## Author contributions

JGB: Project conceptualization, Methodology, Analysis, Writing—Original Draft, Review and Editing; BJM: Methodology, Analysis, Writing—Review and Editing; EK: Analysis, Writing—Review and Editing; CM: Methodology, Analysis, Writing—Review and Editing. All authors contributed critically to drafts and gave final approval for publication.

## Acknowledgments

Support for production of this manuscript was provided by Environmental Solutions & Innovations, Inc.

## Data Availability

All data are from publicly available sources. R scripts will be archived in Zenodo.

## Footnotes

1 https://www.cnn.com/2025/07/26/climate/georgia-bucees-interstate-lights-sea-turtles

