## Supplementary Materials and Code for "Biological Footprint of Artificial Light at Night in Rural and Developing Areas"

Boyles et al.

March 2026

#### Contents

**Physics Equations and Light Behavior** **1**

Omnidirectional Throw . . . 1

Reflected Light . . . 4

**Light behavior with Topography and Vegetation** **8**

**Literature Cited** **10**

### Physics Equations and Light Behavior

#### Omnidirectional Throw

The following equations are well known in physics, but are provided here for easy reference to the reader.

An approximate bound for "throw distance" can be calculated by first converting luminous flux ($\Phi$; expressed in lumens [lm]) to luminous intensity ($I$; expressed in candela [cd]) across a given solid angle ($\Omega$).

$$I = \frac{\Phi}{\Omega}$$

For an omnidirectional light where light emits equally in all directions (e.g., an unshielded bulb assuming no shadow effect of the fixture for simplicity):

$$I = \frac{\Phi}{4\pi}$$

And for a directional light with a focused beam (i.e., a beam angle less than 180°):

$$I = \frac{\Phi}{2\pi(1 - (cos\ \frac{\theta}{2}))}$$

Next, the inverse square law can be applied to determine the distance (in meters) at which a light of a given intensity will produce an illuminance ($E$; expressed in lux) of a specific value, assuming no topography or vegetation to block the light and no atmospheric attenuation:

$$d = \sqrt{\frac{I}{E}}$$

It is further possible to numerically solve for the effect of atmospheric attenuation in calculations of throw distance:

$$E(d) = \frac{I}{d^2} \cdot e^{-\beta \cdot d}$$

Where $E(d)$ is illuminance at distance $d$ (in m) and $\beta$ is the atmospheric attenuation coefficient (in $m^{-1}$). Low $\beta$ values (0.0001 - 0.0005) are typical of very clear air in dry areas like a desert or high mountains (visibility > ~7.8 to 39 km), while values of 0.0005 - 0.001 might be more typical of clear days in rural or suburban areas (visibility from 3.9 to 7.8 km). High values (> 0.01) are typical of heavy fog (visibility < 0.39 km). We use $\beta = 0.0005$ in examples herein. For simplicity, we also ignore "falloff", the tendency for focused lights to be brightest in the center of the beam and weaker progressing toward the edges.

```r
# Necessary libraries
library(ggplot2)
library(dplyr)
library(cowplot)
library(tidyr)

## Input Settings (these can be adjusted)
# range of luminous flux:
lumens_seq <- seq(500, 100000, by = 100)
# linear spacing:
lux_seq <- signif(seq(0.001, 0.25, length.out = 300), 4)
# visibility in km. 7.8 represents clear, rural air:
visibility_km <- 7.8
# attenuation coefficient based on Koschmieder's law:
atmospheric_k <- round(3.912/(visibility_km*1000), 4)
```

##### Omnidirectional Throw in Parameter Space

Now a simple function to calculate throw distance based on flux and illuminance. After defining the function, let's apply across parameter space and plot.

```r
# Throw Function
calculate_omni_throw <- function(lumens, target_lux, k = 0) {
  solid_angle <- 4 * pi
  intensity_cd <- lumens / solid_angle

  if (k == 0) {
    d <- sqrt(intensity_cd / target_lux)
  } else {
    f <- function(d) {
      (intensity_cd / d^2) * exp(-k * d) - target_lux
    }
    d <- tryCatch(uniroot(f, lower = 0.1, upper = 10000)$root,
                  error = function(e) NA)
  }
  return(d)
}

# Calculate values across parameter space
grid_omni <- expand.grid(Lumens = lumens_seq, TargetLux = lux_seq)
grid_omni$ThrowDistance <- mapply(
```

```r
  calculate_omni_throw,
  lumens = grid_omni$Lumens,
  target_lux = grid_omni$TargetLux,
  MoreArgs = list(k = atmospheric_k)
)
```

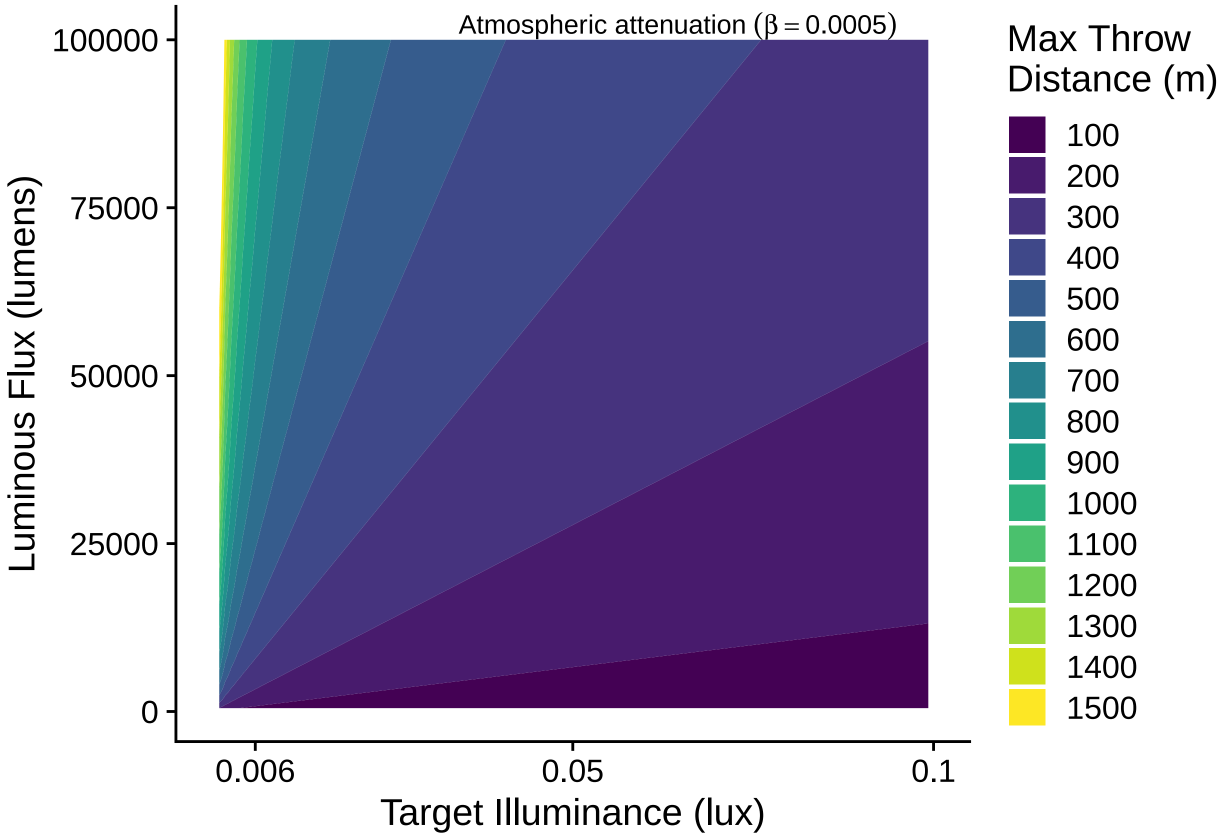


*Figure S-1. The maximum throw distance (in meters) at which a light of a given brightness can illuminate an object to a given intensity. Target illuminance of 0.006 lux approximates starlight on an otherwise dark night (~0.001 - 0.01 lux) while 0.1 lux approximates the low end of expected light around a full moon (but not the brightest possible). Values are grouped into 100 m bins for visual clarity.*

###### Directional Throw in Parameter Space

As described in the main text, throw distance is also augmented with directional lighting and shielding. The calculation operates similarly to the simple omnidirectional light above, using the inverse square law, but with target illuminance held constant (at 0.006 lux, where some animals may be affected by light; Aulsebrook et al. 2022) and the effect of beam angle driving how far throw is possible.

```r
# hold constant at minimum biological effect:
target_lux <- 0.006
# range of luminous flux:
lumens_seq <- seq(1000, 100000, by = 100)
# beam angle, from narrow focused light to wide-angle lighting"
beam_angles_seq <- seq(20, 180, by = 1)
```

```r
# visibility in km. 7.8 represents clear, rural air:
visibility_km <- 7.8
# attenuation coefficient:
atmospheric_k <- 3.912/(visibility_km*1000)
```

```r
# Directional Throw Function
calculate_throw_distance <- function(lumens, beam_angle_deg, target_lux = 0.006, k = 0) {
  half_angle_rad <- (beam_angle_deg / 2) * pi / 180
  solid_angle <- 2 * pi * (1 - cos(half_angle_rad))
  intensity_cd <- lumens / solid_angle

  if (k == 0) {
    d <- sqrt(intensity_cd / target_lux)
  } else {
    effective_illuminance <- function(d) {
      (intensity_cd / d^2) * exp(-k * d) - target_lux
    }
    d <- uniroot(effective_illuminance, lower = 0.1, upper = 10000)$root
  }
  return(d)
}

# Calculate values across parameter space
grid <- expand.grid(Lumens = lumens_seq, BeamAngle = beam_angles_seq)
grid$ThrowDistance <- mapply(calculate_throw_distance,
                             lumens = grid$Lumens,
                             beam_angle_deg = grid$BeamAngle,
                             MoreArgs = list(target_lux = target_lux, k = atmospheric_k))
```

#### Reflected Light

Light reflected into the environment is dependent on not only the characteristics of the light, but also the surface off which it reflects. Specifically, surface characteristics determine the amount of reflectance (albedo; $\rho$). The distance at which the reflectance is equal to the target illuminance ($E$) can be calculated by first calculating the light reflected off the ground and other surfaces (assuming Lambertian reflectance):

$$L = \frac{\Phi \cdot \rho}{A\pi}$$

Where L is the reflected luminance (in cd m$^{-2}$) and A is the area illuminated. As with direct throw of light described above, the inverse square law can be used to calculate the distance at which a reflected light will equal an illuminance ($E$; expressed in lux) of a specific value, assuming no topography or vegetation to block the light and no atmospheric attenuation:

$$d = \sqrt{\frac{L \cdot A}{E}}$$

Notice that area cancels out in these equations if it is assumed the observer can see the entire reflected area, which might be the case for an aerial insect or bat flying above the light. At a lower angle, the observed area decreases.

Finally, it is further possible to numerically solve for the effect of atmospheric attenuation in these calculations:

$$E(d) = \frac{L \cdot A}{d^2} \cdot e^{-\beta d}$$

Now let's see how light behaves in parameter space. We can look at individual lights and a broad spectrum of numbers of lights and light brightness and focus the distance at which specific configurations have biological effects.

##### Reflectance of an Individual Light

As noted in the text, reflected light can reach significant distances, even when shielded. The calculation expands on simple light throw by assuming all environmental surroundings act as Lambertian reflectors. Although the assumption is likely violated in most any real-world application, it allows for estimation of a maximal effect, particularly for volant nocturnal animals.

```r
# Function for calculations
throw_distance_reflected <- function(height_m, lumens, beam_angle_deg, albedo = 0.25,
                                     required_illuminance = 0.006, visibility_km = 23) {
  half_angle_rad <- (beam_angle_deg / 2) * pi / 180
  r <- height_m * tan(half_angle_rad)
  area_m2 <- pi * r^2
  E_ground <- lumens / area_m2
  L_cd_m2 <- (E_ground * albedo) / pi
  # attenuation coefficient based on Koschmieder's law:
  atmospheric_k <- round(3.912/(visibility_km*1000), 4)
  # full illuminated patch:
  A_observed <- area_m2

  f <- function(d) {
    (L_cd_m2 * A_observed / d^2) * exp(-atmospheric_k * d) - required_illuminance
  }

  throw_distance_m <- tryCatch({
    uniroot(f, lower = 1, upper = 10000)$root
  }, error = function(e) NA_real_)

  return(tibble::tibble(
    throw_distance_m = throw_distance_m,
    area_m2 = area_m2,
    E_ground = E_ground,
    L_cd_m2 = L_cd_m2
  ))
}


# Parameter grid: varying height and beam angle
param_grid <- expand.grid(
  height_m = 5,
  beam_angle = 150,
  lumens = seq(1000, 100000, by = 1000),
  visibility_km = seq(.5, 23, by = .5)
)

# Compute throw distances
grid_reflect <- param_grid %>%
  rowwise() %>%
  mutate(
```

```
    throw_distance_m = throw_distance_reflected(height_m, lumens, beam_angle,
                                     albedo = 0.25,
                                     required_illuminance = 0.006,
                                     visibility_km = visibility_km)
  ) %>%
  ungroup()
```

##### Reflectance of Multiple Lights

The same can be done for multiple lights in an array.

```
# Function for calculations
throw_distance_reflected_multi <- function(height_m, lumens_per_light, beam_angle_deg,
                                          albedo = 0.25, required_illuminance = 0.006,
                                          visibility_km, n_lights = 1,
                                          overlap_fraction = 0.3) {
  half_angle_rad <- (beam_angle_deg / 2) * pi / 180
  r <- height_m * tan(half_angle_rad)
  area_single <- pi * r^2

  # 2D area model
  area_total <- area_single * (1 + (n_lights - 1) * (1 - overlap_fraction)^2)

  total_lumens <- n_lights * lumens_per_light
  E_ground <- total_lumens / area_total
  L_cd_m2 <- (E_ground * albedo) / pi

  A_observed <- area_total
  beta <- 3.912 / (visibility_km * 1000)

  f <- function(d) {
    (L_cd_m2 * A_observed / d^2) * exp(-beta * d) - required_illuminance
  }

  d_solution <- tryCatch({
    uniroot(f, lower = 1, upper = 10000)$root
  }, error = function(e) {
    NA_real_
  })

  return(list(
    throw_distance_m_multi = d_solution,
    area_m2 = area_total,
    E_ground = E_ground,
    L_cd_m2 = L_cd_m2
  ))
}


# Parameter grid: varying height and beam angle
param_grid_multi <- expand.grid(
  height_m = 5,
  beam_angle = 150,
```

```r
  lumens = seq(1000, 100000, by = 1000),
  visibility_km = 7.8,
  n_lights = seq(1, 25, by = 1)
)

# Compute throw distances
grid_reflect_multi <- param_grid_multi %>%
  rowwise() %>%
  mutate(results = list(throw_distance_reflected_multi(
    height_m, lumens, beam_angle,
    albedo = 0.25,
    required_illuminance = 0.006,
    visibility_km = visibility_km,
    n_lights = n_lights,
    overlap_fraction = 0.3
  ))) %>%
  unnest_wider(results) %>%
  ungroup()
```

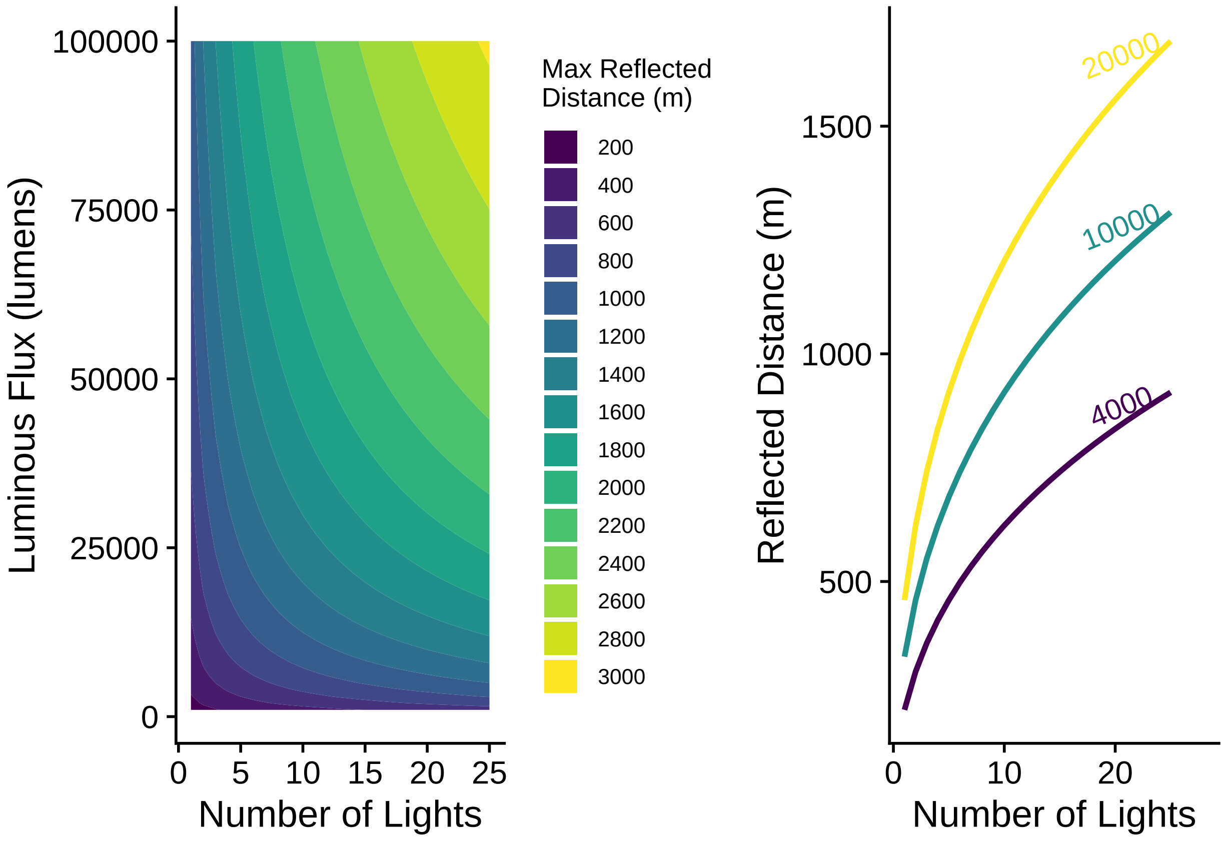


Figure S-2. Effect of multiple lights in an array on throw distance of reflected light into the environment. This simulation assumes an albedo ($\rho$) of 0.25 and a target illuminance ($E$) of 0.006. Asphalt (tarmac) is closer to $\rho = 0.1$, while lightly colored cars or unfinished metal can have $\rho > 0.5$. Left panel: estimated distance of biological effects for light arrays comprised of lights of a given intensity. Right panel: distance of biological effects for lights of three sizes (4000, 10000, and 20000 lumens).

### Light behavior with Topography and Vegetation

Forest height, canopy cover, and elevation are easily transferable to light propagation modeling. Canopy density can be estimated from canopy bulk density (CBD; the amount of burnable fuel in the canopy given in kg m-3) presented by LANDFIRE to an optical cross-section useful in estimating a light extinction rate (k) in the classic Beer-Lambert Law. Here, we used a simple linear scaling factor (effective optical cross-section per unit mass; $\sigma_m$) to accomplish this:

$$k = \sigma_m \cdot CBD$$

There are several lines of reasoning useful in determining the value of $\sigma_m$ (specific leaf area, radiative transfer in vegetation models, forest transmissivity models, etc), and all return values of approximately 2-25, depending on forest type. As a general heuristic, specific leaf area (when expressed as $m^2/kg$) provides a reasonable estimate of $\sigma_m$ although it is not fully analogous. LANDFIRE further includes a vegetation type data layer (LANDFIRE 2024), which has designations for conifer, mixed conifer-hardwood, and hardwood forests. We set $\sigma_m$ to 6, 9, and 12 for the three forest types, respectively.

The manuscript body provides a contextualized overview of this study. The following figures provide aerial imagery alongside model output and topographic relief to demonstrate how light behaves under different environmental conditions. Randomly chosen 50- by 50-km squares were drawn across the U.S. to capture topographic and vegetative variation needed to characterize model (and thus light) behavior.

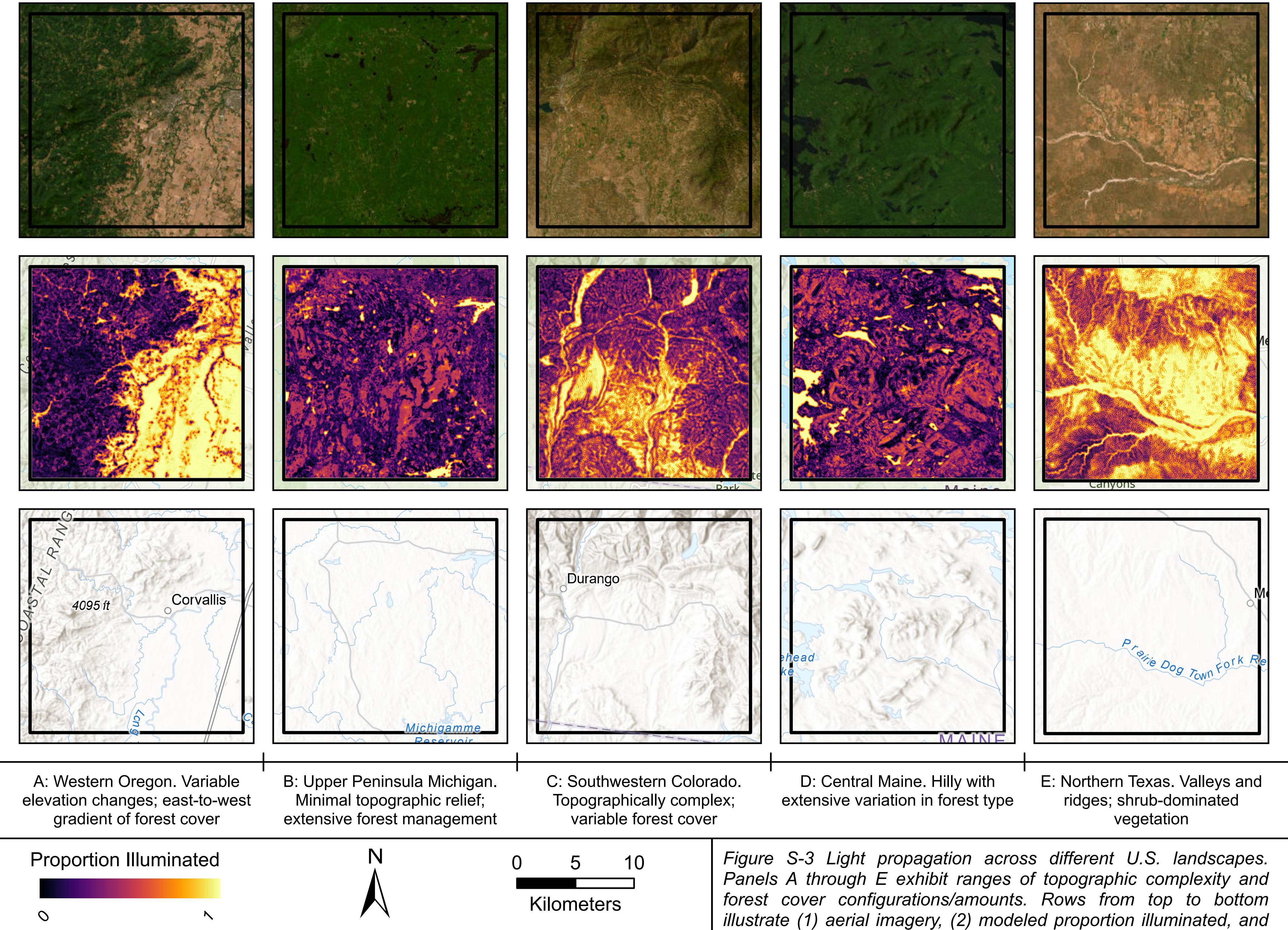


Service Credits: Earthstar Geographics, Esri, CGIAR, USGS, Sources: Esri, TomTom, Garmin, FAO, NOAA, USGS, (c) OpenStreetMap contributors, and the GIS User Community, Sources: Esri, TomTom, Garmin, FAO, NOAA, USGS, © OpenStreetMap contributors, and the GIS User Community, Esri, USGS

*Figure S-3 Light propagation across different U.S. landscapes. Panels A through E exhibit ranges of topographic complexity and forest cover configurations/amounts. Rows from top to bottom illustrate (1) aerial imagery, (2) modeled proportion illuminated, and (3) topographic relief. Generated in ArcGIS Pro vers. 3.5.2. using model outputs from R in (2).*

LANDFIRE. 2024. Fuel Vegetation Type Layer.in U.S. Department of the Interior, Geological Survey, and U.S. Department of Agriculture, editors.
